# The thermotolerant Arabian killifish, *Aphanius dispar*, as a novel infection model for human fungal pathogens

**DOI:** 10.1101/2024.10.08.617174

**Authors:** Rashid Minhas, Tina Bedekovic, Teigan Veale, Larissa L. H. John, Atyaf Saied Hamied, Elliott Scrase, Sienna Read, Chantelle Davies, Hugh Gifford, Rhys Farrer, Steven Bates, Alexandra C. Brand, Mark Ramsdale, Tetsuhiro Kudoh

**Author notes:** These authors equally contributed to this work (Rashid Minhas, Tina Bedekovic). **Author contributions** Conceptualization: TK and MR. Investigation and Methodology: RM, TB, TV, LJ, AH, CD, ES, SR, HG, and TK. Formal analysis: RM, TB, MR, TK. Funding acquisition: MR, RF, SB, AB, TK. Writing - original draft: RM, TB, MR, TK. Writing - review and editing: RM, TB, AB, SB, TK. **Competing interests** The authors declare that they have no competing interests.

## Abstract

*Candida albicans*: a fungal pathogen, can cause superficial and fatal infections in humans. An important virulence factor in *C. albicans* dissemination is the transformation from yeast to an invasive hyphal form, which is favoured at human body temperature. Zebrafish, a useful model for studying *C. albicans* infections, cannot survive at 37°C. Arabian killifish, *Aphanius dispar*, an emerging teleost model can tolerate temperatures up to 40 °C for up to 12 days (independent feeding time) allowing for longer analysis compared to zebrafish. This study introduces *A. dispar* as a thermo-relevant and a more accurate reporter of the virulence mechanisms relevant to *C. albicans* as a human pathogen. Using *A. dispar*, we tested virulence at human skin (30 °C), body temperature (37 °C) and a high fever condition (40°C). Infection by *C. albicans* at 37°C and 40°C significantly increased virulence, reduced survival of AKF embryos and formed invasive hyphal network compared to 30 °C. Two mutant strains of *C. albicans. pmr1Δ* (with aberrant cell surface glycans) exhibited reduced virulence at 37°C, whereas *rsr1*Δ (lacking a cell polarity marker) showed less virulence at 30 °C. Additionally, anti-fungal treatment rescued AKF survival in a dose-dependent manner, indicating AKF’s potential for *in vivo* drug testing. Our data indicates the quantitative and qualitative importance of examining virulence traits at physiologically relevant temperatures and demonstrates an equivalence to findings for systemic infection derived in mouse models. The *A. dispar* embryo therefore provides an excellent *in vivo* model system for assessing virulence, drug-testing, and real-time imaging of host-pathogen interactions.

**Significance Statement:** The virulence of many pathogens is dependent on host temperature. We demonstrate that the *A. dispar* embryo provides an excellent new thermo-relevant alternative to zebrafish and mouse models, which have limitations in terms of the range of temperatures that can be assessed in real-time. In this study, we have assessed *C. albicans* temperature-based virulence, focusing on human body and human skin temperatures (37, 40 and 30 °C, respectively) by examining different genetic backgrounds of *C. albicans* strains. The results indicate different *C. albicans* strains with genetic background show varied virulence depending on temperature indicating importance of examination of virulence mechanisms at physiological temperatures.

## Introduction

*Candida* species are opportunistic fungal pathogens of humans that reside as commensals in the oral, gastrointestinal, and vaginal tracts of healthy individuals (Brown et al., 2012; Lewis and Williams, 2017; Zeise et al., 2021). They cause infections of the skin, mucosa and nails (MacCallum, 2010) which can become chronic (Rautemaa-Richardson et al., 2022), and can cause vaginal infections, which in some cases are recurrent (Sobel et al., 1998). For immunocompromised patients, such as those going through chemotherapy, immune suppression or organ transplants, infection can become disseminated, causing invasive candidiasis (Brown et al., 2012; Maksymiuk et al., 1984), often associated with high mortality rates (∼45%) (MacCallum, 2010). Candidiasis is a major problem in hospitals (Markogiannakis et al., 2009; Wisplinghoff et al., 2006) as *Candida* species can adhere to, and form biofilms on, biotic and abiotic surfaces, including medical implants (Ramage et al., 2006). *Candida albicans* alone is estimated to cause over 150 million mucosal infections and roughly 200,000 deaths every year (Richardson, 2022). *C. albicans,* the most prevalent species, is polymorphic and able to switch from yeast to filamentous hyphae in the presence of serum, high CO_2_ levels and at temperatures above ∼35 °C. Hypha formation is an important virulence factor, promoting tissue invasion and damage, biofilm formation and inflammation through secretion of host damaging proteinases, as well as changes in the cell wall that help to evade the host immune system (Baillie and Douglas, 1999; Noble et al., 2017). Virulence traits can be examined *in vitro*, but to truly understand host-pathogen interactions, particularly in real time, *in vivo* models are crucial.

Numerous organisms have been exploited as models in which to study *C. albicans*, all offering different advantages and drawbacks. The murine model has been used to model gastrointestinal (de Repentigny et al., 1992), vaginal (Sobel et al., 1985), oropharyngeal (Solis et al., 2018) and systemic candidiasis (Hebecker et al., 2016). Various mouse models of *C. albicans* infection have been developed including an intravenous challenge model to study invasive candidiasis using immunocompetent mice, oral cavity infection in immunocompromised mice and skin infection to study systemic immunity (Clancy et al., 2009; Fajardo et al., 2021; Kashem et al., 2015; Seiser et al., 2024). Due to ethical concerns surrounding the use of mammals, in addition to high maintenance costs and limited sample sizes, alternatives would be welcome. Over a decade ago zebrafish embryos have become a popular model for *Candida* infection studies (Chao et al., 2010; Rosowski et al., 2018; Seman et al., 2018). Many transgenic zebrafish lines are now available, including those with altered immunity, which aid the study of *C. albicans* virulence factors and host-pathogen interactions (Torraca et al., 2014). Zebrafish embryos can also be used to directly study the immune cell-pathogen interaction where macrophages have been found to inhibit the yeast-hypha transition and other key virulence traits *in vivo*, implicating reactive oxygen and nitrogen species (Brothers et al., 2011). Zebrafish eggs and embryos are transparent, allowing non-invasive real-time imaging of fungal growth behaviour in infected embryos (Brothers and Wheeler, 2012). Images of yeast cells and hyphae present in the tail, following hindbrain injections, have confirmed that zebrafish can be used to study disseminated infection and follow the growth and invasion of the fungal cells for up to 5 days (Brothers et al., 2011). In the zebrafish model of candidiasis, the Wheeler group has demonstrated the intravital imaging of yeast cells whilst modulating the temperatures between 21 °C and 33 °C (Seman et al., 2018). However, the optimal temperature for zebrafish development is 28 °C and, being unable to survive at 37 °C, they cannot serve as a fully representative model of human pathogenesis (Kimmel et al., 1995). This is particularly relevant when studying the virulence of fungal strains like clinical isolates and mutant strains (like *pmr1Δ*) of *C. albicans*. Medaka are another small fish that are an important model for studying infections and can survive at high temperatures (Broussard and Ennis, 2007) however, due to the rhythmic contraction of the yolk and the hairy chorion, they are not easily used for time-lapse analyses or for high-resolution, high-throughput *in vivo* imaging. Invertebrate models, such as *Drosophila melanogaster*, *Galleria mellonella*, and *Caenorhabditis elegans*, have also been employed to study candidiasis (Davis et al., 2011; Fuchs et al., 2010; Jacobsen, 2014; Pukkila-Worley et al., 2011). In contrast to other invertebrate models *G. mellonella* larvae can be maintained at temperatures up to 37 °C (Serrano et al., 2023). However, due to their significant evolutionary distance and physiological differences from mammals, there is a growing need for models with a closer evolutionary relationship, adaptive immune system and more similar internal organs.

Arabian killifish, *Aphanius dispar*, are euryhaline thermotolerant teleost fish ubiquitous to coastal waters surrounding the Arabian Peninsula, the Red Sea, and the Eastern Mediterranean Sea. Adults and embryos have been observed naturally in a wide range of temperatures, including 30-37 °C. Despite their largely herbivorous tendencies, *A. dispar* fish are larvivorous and therefore have undergone successful trials as biological control agents of mosquito larvae (Frenkel and Goren, 2000; Haas, 1982). Arabian killifish embryos have also been used as a model for assessing the risk of marine pollution to the Arabian Gulf coastal waters (Saeed et al., 2015). *A. dispar* embryos have a transparent yolk and chorion (Alsakran et al., 2024), which permits high-resolution live imaging of infection dynamics. Moreover, their small size and lack of rhythmic contraction make this species an ideal model for both high throughput and high-resolution real-time imaging (Hamied et al., 2020). Here we show that Arabian killifish embryos have several features that offer the potential to accelerate research into *C. albicans* infection and lead to the development of new and more effective treatments for candidiasis and other fungal infections.

## Results

### Use of the Arabian killifish model demonstrates that *C. albicans* virulence is enhanced at 37 and 40 °C compared to 30 °C

To study the significance of temperature in the level of virulence *in vivo*, *C. albicans* cells were introduced by microinjection into the yolk of *A. dispar* embryos at 3 dpf and incubated at typical human skin temperature (30 °C) and human body temperature (37 °C). Embryos injected with *C. albicans,* and the PBS and un-injected controls, were transferred to a 96-well plate and imaged by the ACQUIFER multi-well imaging system (Fig. 1A). To monitor the heartbeat as a marker for live embryos, 5 successive images were captured, and time-of-death of injected embryos was recorded when no heartbeat was observed (Fig. 1B). Our data using two different *C. albicans* control strains (NGY152 and ACT1p-GFP) showed that mortality followed infection more quickly when embryos were incubated at human body temperature (37 °C) compared to human skin temperature (30 °C) (Fig. 1C). At both temperatures, the majority of infected embryos died within 3 d, but the MTD (mean time to death) for NGY152 at 30 and 37 °C were ∼62.5 hpi and ∼32 hpi, respectively. While the MTD for the ACT1p-GFP strain at 30 and 37 °C were 63.5 hpi and 25 hpi, respectively. Overall, the MTD in both control strains are closely matched. In both strains, there was a significant acceleration in death rate when embryos were incubated at normal human body temperatures. The death rate of the control strain NGY152 was also similar to that of the wild-type parent strain, SC5314 (Fig. S1A). As observed at 37 °C, a similar mortality profile was observed at the pyrexic (fever-related) temperature of 40 °C (Fig. S1B). To support and validate the use of heartbeat arrest as a marker for embryo death, the reduction of yolk diameter was also measured. A few hours after cardiac arrest in the infected embryo, the yolk ball started to shrink and reduced in diameter, with a horizontal reduction of 30 % and vertical reduction of 36 % by 41.5 hpi) (Fig. 1D). The MTD, determined by the initiation of yolk shrinkage, for 30 °C and 37 °C were ∼62 hpi and ∼38 hpi, respectively, for the NGY152 control strain. This method confirmed that accelerated lethality occurred at 37 °C compared to 30 °C (Fig. 1E).

**Fig. 1.**
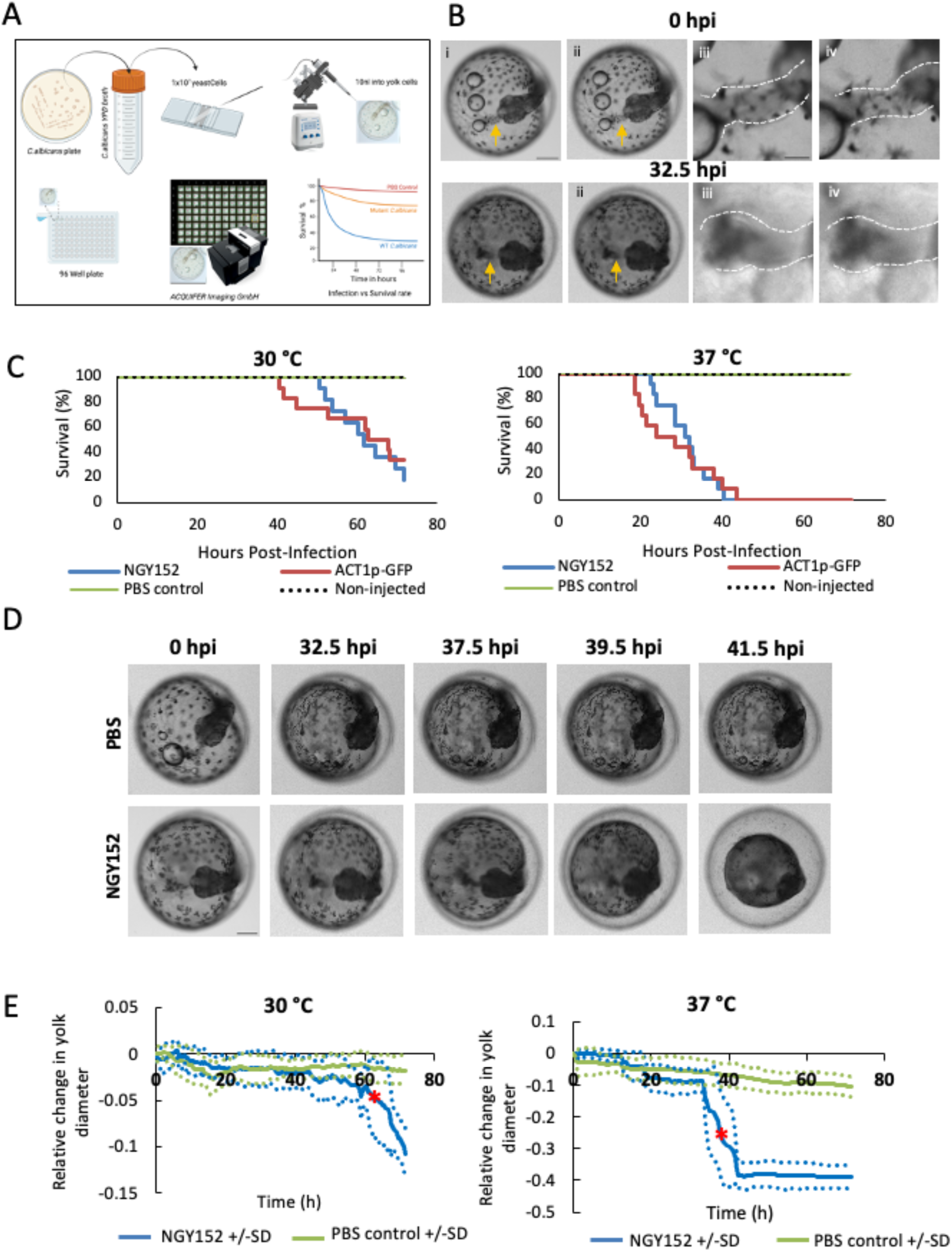
C. albicans induces earlier and higher lethality of the A. dispar embryo at 37 °C compared with 30 °C. (A) Workflow schematic showing *C. albicans* infection studies in *A. dispar* embryos, with monitoring of the infected embryos using ACQUIFER and survival rate analysis. (B) Representative brightfield images of a NGY152-infected WT *A. dispar* embryo maintained at 37 °C. 0 hpi (top panel), and 32.5 hpi (bottom panel) (i and ii in each panel, scale bar = 1000 μm) taken using the ACQUIFER Imaging system at 4X magnification and two zoomed in images indicating the heartbeat arrest (iii and iv in each panel, scale bar = 250 μm). Yellow arrows show the heart region and white dotted lines outline the area of the cardiac tube. (C) Survival percentages of NGY152, *ACT1p-GFP*, PBS control and non-injected WT *A. dispar* embryos over a 72-hour period maintained at 30 °C (left panel), or 37 °C (right panel). The survival experiment was performed in triplicate (n=3) with 17 embryos per group. Plotted values represent the mean at each timepoint. (D) Representative brightfield images were taken at 4X magnification by the ACQUIFER Imaging system of a PBS-injected (top panel) and NGY152-infected (bottom panel) WT *A. dispar* embryo maintained at 37 °C for 0, 32.5, 37.5, 39.5, and 41.5 hours post infection (hpi). Death of the NGY152-infected embryo occurred at ∼32.5 hpi whereby the yolk mass starts to decrease. (E) Average relative change in yolk diameter for NGY152- and PBS-injected WT *A. dispar* embryos maintained at either 30 °C (left panel) or 37 °C (right panel) over a 72-hour period. An average of vertical and horizontal measurements for each embryo were taken, accounting for embryo rotation, the non-spherical nature of the yolk and nonuniform yolk shrinkage. Plotted values represent the mean of 2 embryos (each represented as dotted lines) of each timepoint with standard deviation fit to error bars. Red asterisk denotes the MTD.

### *C. albicans* proliferates more rapidly *in vivo* at 37 °C compared to 30 °C

To examine the differential growth responses of *C. albicans* cells in embryos at the two host-relevant temperatures (30 °C and 37 °C), infected *C. albicans* cells were recovered from individual embryos at 5 time-points post infection and cell number analysed using a colony forming unit (CFU) assay (Fig. 2A). CFU counts were significantly higher at 24 h post infection in populations taken from embryos incubated at 37 °C compared with 30 °C – despite the known issues of using CFU counts for fungi that produce mixed cell types (yeast-pseudohyphae, hyphae). To further assess the total biomass within embryos, a tdTomato labelled *C. albicans* strain was injected into embryos and the quantity of the tdTomato protein level was determined using Western blotting (Fig. 2B). The data showed a similar rapid growth of tdTomato labelled *C. albicans* by 24 hpi, which increased further by 48hpi (Fig.2B).

**Fig. 2.**
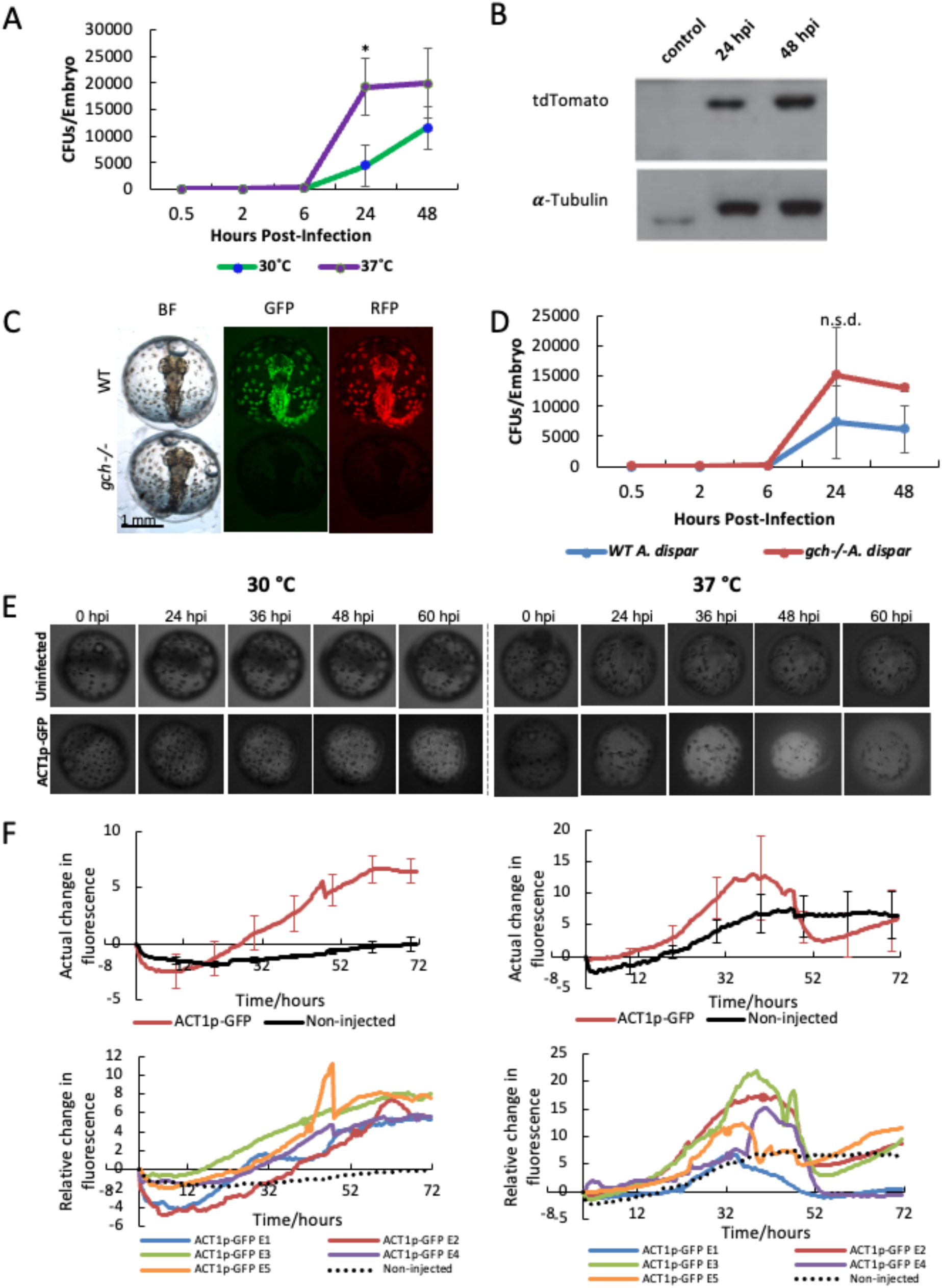
Growth of the C. albicans is accelerated at 37 °C as compared to 30 °C and remain consistent in WT and mutant A. dispar strains. (A) Fungal burden of SC5314 of *C. albicans* strain was determined per embryo by homogenising and plating 5 embryos separately at each time point (0.5, 2, 6, 24 and 48 hpi) and compared at 30 °C and 37 °C. Three independent experiments (n = 3) were done and datapoints represent mean with error bars. Asterisk represents significant difference at the 24 hpi (p<0.05). (B) Representative western blot showing the time-course of fungal infection (1×10^8^ cells/ml CAF2-dTomato) in the *A. dispar* embryos up to 72 hours. Proteins were detected utilising the mCherry antibody (Abcam) and a *α*-Tubulin antibody used as a loading control. (C) WT and *gch-/-* embryos at 3 dpf imaged with transmitted/bright field (BF), GFP filter, and RFP filter (Hamid et al. 2020). *gch-/-* line was subsequently used for ACT1p-GFP. (D) WT and *gch-/- A. dispar* embryos were injected with 5 x 10^6^ cells/ml of ACT1p-GFP *C. albicans* suspension into the yolk at 3 dpf and incubated at 37 °C for a maximum of 48 hours. At each time point, 3-5 embryos were separately homogenised to enumerate the colony forming units (CFUs) after plating. Three independent experiments (n = 3) were done and datapoints represent mean with error bars (n.s.d. = non-significant difference) (E) Fluorescent images taken at 4X magnification, wavelength 470 nm, by the ACQUIFER Imaging Machine of a un-injected (top panel) and ACT1p-GFP-injected (bottom panel) *gch(-/-) A. dispar* embryos at 30 °C and 37 °C at 0, 24, 36, 48 and 60 hpi. (F) Average relative change in fluorescence for ACT1p-GFP- and non-injected *gch(-/-) A. dispar* embryos maintained at either 30 °C (left panel) and 37 °C (right panel) over a 72 hr period. Relative change in fluorescence was standardised using background fluorescence. Plotted values represent the average of 5 embryos at each timepoint with standard deviation fit to error bars (bottom panels).

To facilitate embryo imaging, we previously reported the development of a GTP cyclohydrolase (*gch*) mutant line of *A. dispar,* which lacks the fluoroleucophores that generate autofluorescent interference (Fig. 2C) (Hamied et al., 2020). To assess if deletion of *gch* alters the immune response of *A. dispar* embryos to *C. albicans*, the fungal burdens infected WT and *gch-/-* killifish embryos were determined at 5 time points up to 48 hpi (Fig. 2D). The CFU/embryo followed a similar trend in WT and *gch-/-* embryos with no significant difference between both fish lines at any time point (Fig. 2D). The fungal burden started to increase at 6 hpi and reached its maximum at 24 hpi, with similar CFUs at 48 hpi. To monitor real time growth of *C. albicans in vivo*, we also examined the growth of fluorescent *C. albicans* (ACT1p-GFP) by quantitatively measuring the fluorescence in the non-fluorescent *gch(-/-)* infected embryos. The embryos were incubated at 30 °C or 37 °C in the ACQUIFER imaging system and the level of fluorescence in the infected embryo was quantified every 30 min. The infected embryos showed constant elevation of fluorescence after infection compared to controls and reached the maximum level of fluorescence at around 60 hpi and 36 hpi at 30 °C and 37 °C, respectively (Fig. 2E). These time points approximately aligned with the time of MTD for each temperature, suggesting the growth of *C. albicans* continues until the death of the host (Fig. 2F). To examine the morphological transformation of *C. albicans* in the *A. dispar* embryos in detail, fluorescently labelled *C. albicans* (Eno1-iRFP) were injected onto the surface of the yolk, incubated at 37 °C and examined by time lapse imaging. At 20 min post-infection, development of transition of yeast to hyphae was observed in some *C. albicans* cells (Fig. 3B). Subsequently, these elongated and formed a dense network of filamentous cells by 24 hpi (Fig. 3C).

**Fig. 3.**
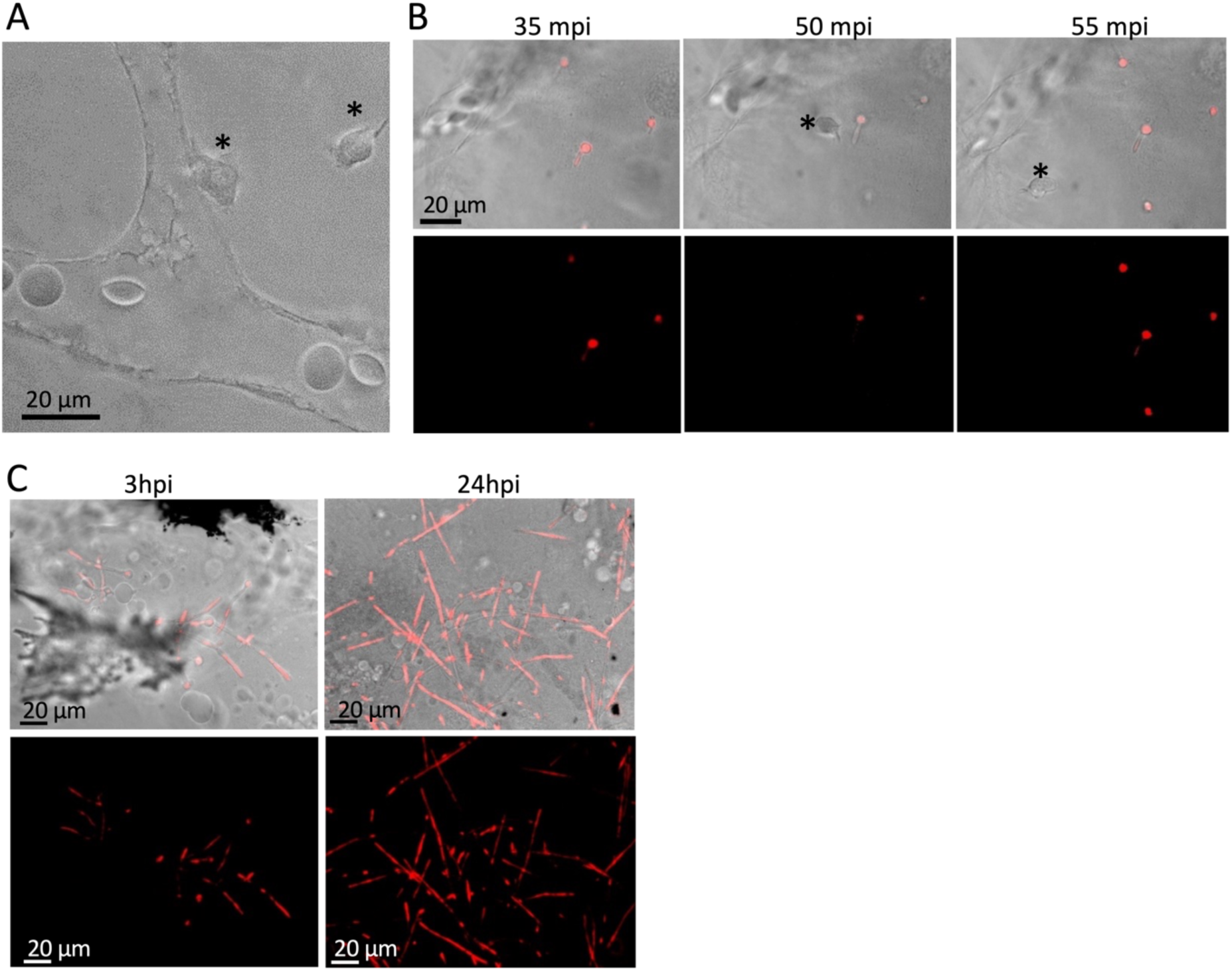
C. *albicans* undergoes morphogenesis in the yolk of *A. dispar* embryos. (A) Representative image of a blood vessel at the yolk surface in an uninfected embryo. Asterisks indicate immune cells. (B) Representative images at 3 time points in the morphological transition of *C. albicans* expressing Eno1-iRFP injected near the yolk surface of the *gch-/-* line. Asterisks indicate immune cells. mpi = min post infection. Top panel: DIC-RFP merged images. Lower panel: images taken using RFP filter. (C) The yolk of dechorionated *gch-/- A. dispar* embryos at 3 dpf were injected with with 1 x 10^8^ cells/ml (right panel) or 5 x 10^7^ cells/ml (left panel) of Eno1-iRFP *C. albicans* suspension, incubated at 37 °C and imaged at 3 hpi and 24 hpi. DIC and RFP fluorescence images were merged. Scale bar = 20 μm.

### *C. albicans rsr1*Δ and *pmr1*Δ mutant strains reveal temperature-dependent differences in virulence compared with the control strain

To examine if *C. albicans* with mutant genetic backgrounds show distinct differences in virulence at various temperatures, we compared the virulence of the WT DAY *C. albicans* strains with *rsr1*Δ at 28 °C (in *D. rerio*), and *rsr1*Δ and *pmr1*Δ at 30 and 37 °C (in *A. dispar*). *Rsr1* encodes a small GTPase involved in the localisation of Cdc42 which plays a role in polarised growth in yeast and hyphal development in *C. albicans.* The *rsr1*Δ mutant has previously been shown to have a virulence defect in a mouse model of infection (i.e., at 37 °C) (Brand et al., 2008; Pulver et al., 2013; Yaar et al., 1997). We infected the yolk sacs of both zebrafish (*D. rerio*) and Arabian killifish embryos with WT and *rsr1*Δ strain and monitored survival for 72 h. In zebrafish at 28 °C, both WT and *rsr1*Δ strains showed similar levels of virulence (Fig. 4A). However, in *A. dispar,* the rsr*1*Δ strain was less virulent at 30 °C. As seen previously, virulence in the control strain was more pronounced at 37 °C than 30 °C, and weak virulence was also observed in the *rsr1*Δ mutant at the higher, but not the lower, temperature (Fig. 4B). This suggests that mutant phenotypes might be missed if not investigated at human body temperature.

**Fig. 4.**
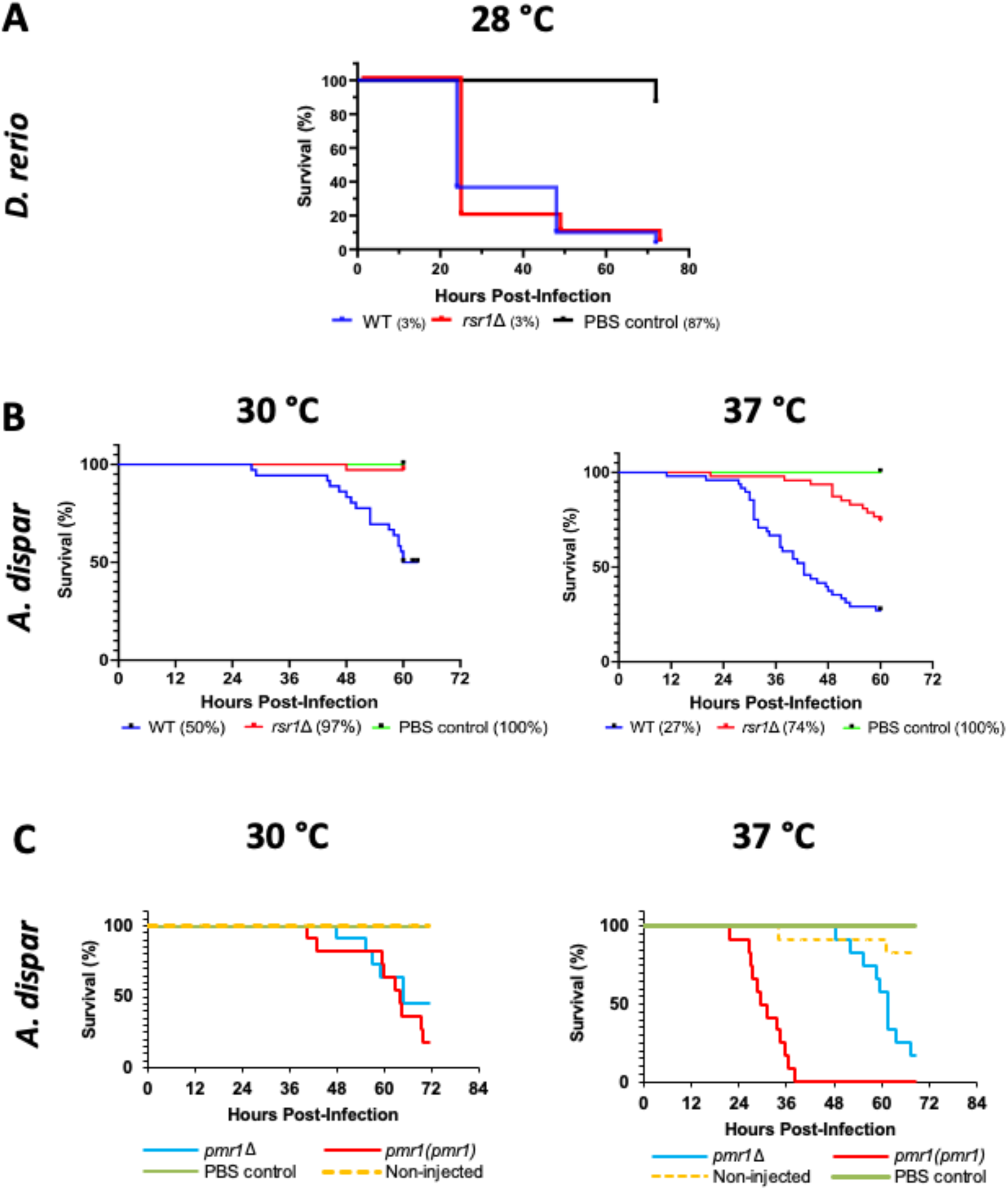
Differential response of C. albicans mutants with attenuated virulence to different temperatures. (A) Survival graphs of Zebrafish (D. rerio) infection incubated at 28 °C with control strain, rsr1Δ mutant strain, and PBS control (n=30 embryos per strain, n=2). (B) Survival plots of A. dispar yolk infection with the WT, rsr1Δ mutant strain, and PBS control incubated at 30 °C (left panel) (3 bio reps, n=12 for each, Log-rank (Mantel-Cox) test = <0.0001) and 37 °C (right panel) (3 bio reps, n=12 for each strain, log-rank (Mantel-Cox) test = <0.0001). (C) Survival plots of A. dispar yolk infected with the pmr1Δ mutant, the pmr1+PMR1 (single copy re-integrant), PBS control, and non-injected embryos incubated at 30 °C (left panel) and 37 °C (right panel) (1 bio rep, n=12).

The Pmr1 symporter is responsible for the import of metal co-factors into the Golgi, where they are required for mannosylation of cell surface proteins. These mannose chains act as potential PAMPS - pathogen-associated molecular patterns (Bates et al., 2005). At 30 °C, *pmr1*Δ showed similar virulence to WT at 30 °C. However, at 37 °C in which WT showed highly elevated virulence but *pmr1*Δ did not show marked increase of virulence (Fig. 4C).

These two examples indicate that genetic background plays an important role in host-pathogen interactions and virulence outcomes in different infection conditions such as 30 °C (e.g. skin infection) and 37 °C (organ infection), highlighting the necessity of undertaking *in vivo* assessment of strains at medically relevant temperatures, as enabled by use of the Arabian killifish embryo model.

### Stage-dependent suppression of the virulence of *C. albicans* by fluconazole

To investigate the sensitivity of the infection model to treatment with antifungals, fluconazole doses were applied to *C. albicans* injected embryos at 2 hpi (4 mg/L to 16 mg/L) or 6 hpi (4 mg/L to 64 mg/L). Although application of the drug reduced the lethality of infection within host embryos in a dose-dependent manner, the outcome was highly time dependent. Early fluconazole treatment at 2 hpi, even at the lowest dose of 4 mg/L, rescued 58 % of infected *A. dispar* embryos, with survival rates increasing to 100 % at 16 mg/L (Fig. 5). However, when the infected *A. dispar* embryos were treated with fluconazole at 6 hpi, higher mortality occurred, even with the highest dose tested of 64 mg/L (Fig. S2). These results demonstrate a time and dose-dependent effect, with increasing fluconazole concentrations directly correlating with enhanced embryo rescue rates, but only if the embryos were treated earlier as shown previously in mice (MacCallum and Odds, 2004). These findings suggest that there is a crucial early infection phase when the host, in combination with the drug, can suppress the growth of *C. albicans*.

**Fig. 5.**
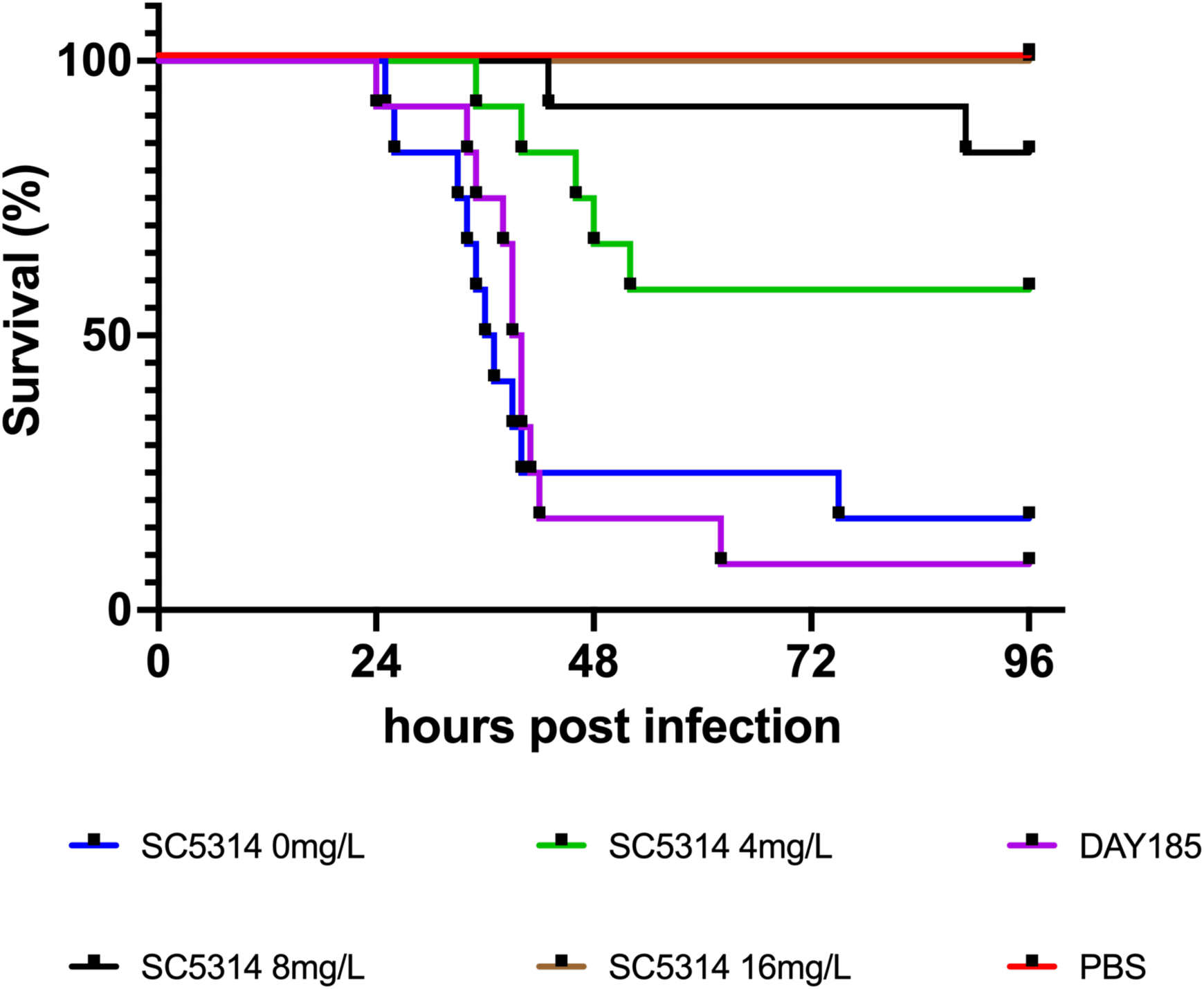
Early intervention with fluconazole is required for successful treatment of C. albicans infection in A. dispar embryos. *A. dispar* embryos were infected at 3 dpf with 5 x 10^6^ cells/ml cells of *C. albicans* (SC5314) and incubated at 37 °C. Embryos were treated at 2 h post infection with 4, 8, or 16 mg/L fluconazole. Survival curves show the effect of fluconazole on *C. albicans* infected *A. dispar* embryos over 96 h. Control groups included PBS alone (no fungal cells), the SC5314 wild-type clinical isolate, and Day185, a control parental strain for many *C. albicans* mutants and included for comparison with SC5314. A statistically significant difference (p-value < 0.0001) was observed between the control and experimental groups, (2 bio reps, n=12 for each condition) (log-rank (Mantel-Cox) test). Please note PBS control and 16 mg/L treatment graph lines overlap.

## Discussion

### Arabian killifish: a novel thermo-relevant and convenient tool kit for fungal infection studies

While murine and existing aquatic embryo host models (zebrafish and medaka) offer valuable tools for studying host-pathogen interactions and virulence, they also present certain limitations. For example, zebrafish embryos cannot survive at human body temperatures including fever conditions such as 40 °C, which can affect the physiology and virulence of the infected pathogens (Segal and Frenkel, 2018). Even though mouse models offer consistency with human body temperature it is challenging to recapitulate fever conditions. Besides, real-time imaging of host-pathogen interactions is not feasible in mice, thus restricting our ability to observe the dynamics of infection progression. While invertebrate models like *D. melanogaster*, *G. mellonella*, and *C. elegans* have been widely used to study human pathogens, their lack of homologous immune systems, and internal organs such as the heart cavity, kidneys, swim bladder, liver, and brain limits their suitability for drug screening and side-effect studies targeting specific organs.

Here, we used a novel *in vivo* thermotolerant host model, *A. dispar* embryos, for studying *C. albicans* infections at human skin (30 °C) and internal body temperature (37 °C). The small size and transparency of the embryos facilitated real-time visualization of fungal growth behaviour and host-*Candida* interactions. Furthermore, *A. dispar* larvae exhibit an extended independent feeding period of 11 days post-fertilization (dpf) compared to 5 dpf for zebrafish larvae, thus the use of *A. dispar* larvae enables the monitoring of experiments over an extended timeframe. The real-time data collection capabilities of optically transparent *A. dispar* embryos eliminates the need for extensive mouse culling at sequential time points, making them a superior alternative to infection studies conducted on mice. Large clutch size and the small size of the killifish embryos also allow the study of post infection mechanisms and disease progression in a 96-well plate format. A further potential advantage of the Arabian killifish embryo model lies in its complex anatomy, which allows for infection through multiple routes in an intact host that harbours a functional innate immune system. Arabian killifish have several additional advantages specifically over zebrafish for some experiments. Firstly, the yolk surface is essentially a 2D flat space where endothelial cells develop dense vasculature network and immune cells are actively migrating in the area, offering an ideal platform for monitoring host-pathogen interactions. As the pathogen and immune cells do not move away from the same Z-axis and therefore can be easily imaged over longer period of time (Neiffer and Stamper, 2009; Owen and Kelsh, 2021). Moreover, at the 3-5 dpf stage, unlike zebrafish embryos, killifish embryos remain immobile within the chorion, facilitating time-lapse imaging with minimum use of anaesthesia. Given that anaesthesia can induce embryonic stress (Matthews and Varga, 2012), reduced use of anaesthesia during live imaging is advantageous. The highly transparent chorion of the Arabian killifish also eliminates the need for chorion removal, enabling high-throughputs imaging with relatively high resolution (Fig. 3). These features facilitate experiments that require real-time observation of embryonic development and interaction with the pathogen and the host immune system.

An early potential drawback of the Arabian killifish embryo was the presence of highly fluorescent pigment cells, fluoroleucophores (Hamied et al., 2020) which interfere with the imaging of fluorescently-labelled *C. albicans* cells. However, we have previously generated a *gch -/-* mutant Arabian killifish line in which the fluorescent pigment in fluoroleucophores is absent, allowing for highly sensitive fluorescent imaging (Fig. 2C). This fish line serves as a valuable tool equivalent to the pigment-deficient casper zebrafish mutant strain, used for imaging experiments (White et al., 2008). Our construction of this mutant also highlights the ease with which molecular genetic tools can be applied to create new Arabian killifish lines to facilitate exploration of the role of key immune components.

### *Candida albicans* and temperature-based virulence

To investigate the influence of temperature on virulence, we employed *pmr1* and *rsr1* mutant strains (Table1) in 3 dpf *A. dispar* embryos and monitored disease progression and embryo survival at human skin temperature (30 °C), body temperature (37 °C) and pyrexic temperature (40 °C). To determine the precise time-course of lethality, an automated imaging system was employed, which recorded mortality every 1-h post-infection. Our data shows an acceleration in lethality at 37 °C (peaking at 2 d post-infection), compared to 30 °C, after 72 hpi (Fig. 4). Our findings suggest a delay of approximately 24 h in mortality at 30 °C, highlighting the significance of using the appropriate temperature for evaluating medically relevant pathogenesis. This observation aligns well with the well-known temperature-driven dimorphism of *C. albicans*, in which invasive hyphae are formed (Fig. 3). However, we did not observe any major change in the development of virulence at the pyrexic temperature. Most embryos died between 24- and 48-h post-infection (hpi) (Fig. S1). In contrast to yeast cells, hyphae exhibit enhanced capabilities for adhering to host cells, penetrating tissues, and evading the immune system (Rooney and Klein, 2002). This potentially explains the disparity in virulence between *C. albicans* at human skin temperature (30 °C) and human body temperature (37 °C) (Dupont, 1995; Ernst, 2000; Martin et al., 2011).

Infection studies using mutant strains *pmr1*Δ (which has a weakened cell wall due to defective surface-protein glycosylation) (Bates et al., 2005) and *rsr1*Δ (which has a hypha formation defect) (Bedekovic et al., 2020) further corroborates the important role in fungal growth and pathogenic behaviour played by human body temperature. There was significant differential virulence between the wild-type strain and the mutants in the two tested temperatures. WT, *pmr1*Δ and *rsr1*Δ all showed distinct pattern of virulence depending on the temperatures suggesting the virulence different genetic background of *C. albicans* can exhibit differential virulence in the skin infection around 30 °C and body infection at around 37 °C. Similar to murine studies, where the *pmr1*Δ mutant exhibited markedly reduced virulence (Bates et al., 2005), with all mice surviving the experimental period, *A. dispar* embryos demonstrate delayed mortality at 37 °C. Notably, the *pmr1*Δ+*PMR1* re-integrant and the *pmr1*Δ mutant exhibited little difference in survival compared to the control strain when incubated at 30 °C. Previous studies have shown that, *rsr1*-deficient strains have reduced hypha formation and agar penetration, that is linked to C. *albicans* virulence (Bedekovic et al., 2020). No differences in *rsr1* viability have been reported at 30 and 37 °C (Pulver et al., 2013). The low virulence of *rsr1*Δ at normal human body temperature (37 °C) is consistent with findings in mouse models (Yaar et al., 1997), indicating that the Arabian killifish model recapitulates findings derived in animal models. Thermal shifts from 30 °C to 37 °C, in combination with 20% serum exposure results in germ tube lysis and cell death of approximately 40% of the *rsr1* cells *in vitro*. Our results indicate the importance of studying morphological transitions *in vivo*, where pathogens may behave differently. The data suggest that infection experiments using animal models at non-human body temperatures may lead to spurious conclusions about the relative virulence of pathogen strains with modified genetic backgrounds.

### Examination of antifungal drug susceptibility using the *A. dispar* embryo

Arabian killifish are also an excellent high throughput model for testing dose and time specific effects of an antifungal drug. When fluconazole was added in water at 2 hpi, the drug effectively suppressed the *C. albicans* infection both in terms of fungal burden and embryo lethality. However, when applied at 6 hpi, fluconazole treatment showed only a subtle effect and could not reduce lethality. These data suggest that the critical time window when fungal growth can be controlled by an antifungal is at a very early stage of infection, that is, at around 2 hpi. Once the initial growth and differentiation of the infected *C. albicans* reaches a level at which colonisation is established, the antifungal can no longer suppress its lethality in the host. This work therefore established a suitable time-course for future drug discover protocols using the Arabian killifish model. In summary, Arabian killifish embryos are an ideal model for testing the efficacy of antifungal drugs at a range of temperatures and could provide valuable clinical insights on the efficacy of drugs at normal and pyrexic conditions.

## Conclusion

Arabian killifish embryos, with their small size, transparency, and extended independent feeding period, offer a unique and valuable *in vivo* model for studying infections at human body temperature. Their real-time imaging capabilities and complex anatomy facilitate detailed observation of host-pathogen interactions and immune system responses. Additionally, the availability of *gch* mutant killifish lines lacking fluorescent pigment cells further enhances the suitability of this model for fluorescent imaging studies and demonstrate the ease with which novel mutants representing disease models can be generated. Here we investigated the use of the Arabian killifish as a promising tool for elucidating the mechanisms and factors influencing *Candida albicans* virulence and pathogenesis at key physiological temperatures with different genetic backgrounds of pathogenetic strains. The model can be further developed to investigate a range of pathogens (fungal and bacterial) where human body temperature is a critical determinant of pathogen physiology.

## Materials and Methods

### *A. dispar* husbandry and egg collection

All animal procedures were approved by the Biosciences Ethics Committee at Faculty of Health and Life Sciences, University of Exeter. *A. dispar* were maintained in the Aquatic Resource Centre (ARC), University of Exeter. Wild-type (WT) and *gch (-/-)* mutants of *A. dispar* (Hamied et al., 2020) were used for fungal infections. *A. dispar* were kept in tanks at 28 °C on a 12 h light: 12 h dark cycle in a recirculation system supplies 35 ppt (35 g/kg) artificial seawater (ASW). Natural spawning was used to collect eggs. Egg chambers were placed in tanks the night before and removed 60 minutes after light was turned on. Eggs were sieved out of chamber water and rinsed in ASW. Eggs were kept, ∼30 per Petri dish in ASW and incubated at 28 °C. All embryos were terminated before hatching with 1 % w/v Virkon solution.

### C. albicans culture

The *C. albicans* strains used in this study are listed in Table 1*. C. albicans* cultures were kept on yeast extract peptone dextrose agar (YPD, 1% w/v yeast extract, 2% w/v peptone, 2% w/v glucose, 2% w/v bacteriological agar) plates at 30 °C. Plates were remade weekly from a single colony and incubated at 30 °C for 48 h before storing at room temperature. Before injection, 10 ml YPD broth was inoculated with a single colony and incubated at 30°C with 180 rpm shaking overnight. The next day, YPD broth was removed, and cells were washed twice with 1X phosphate-buffered saline (PBS) by centrifugation at 13000 rpm for 1 minute. Cells were resuspended in 1 ml 1XPBS and diluted to the appropriate concentration using 10% v/v phenol red (Sigma-Aldrich) in PBS to aid visualisation during injection.

**Table 1:** Candida albicans strains used in this study.

| Strain name | Genotype | Reference |
| --- | --- | --- |
| SC5314 | Clinical isolate | (Gillum et al., 1984) |
| BWP17 | *ura3::λimm434/ura3::_λ imm434 his1::hisG/his1::hisG*<br>*arg4::hisG/arg4::hisG* | (Wilson et al., 1999) |
| CAI4 | *ura3::λimm434/ura3::λ imm434* | (Fonzi and Irwin, 1993) |
| NGY152 | (CAI4) *RPS1/rps1Δ::Clp10* | (Munro et al., 2005) |
| WT-Day185<br>(BWP17+Clp30) | *ura3::imm434/ura3::imm434*<br>*his1::hisG::HIS1/his1::hisG arg4::hisG::ARG4-*<br>*URA3/arg4::hisG* | (Davis et al., 2000) |
| *ACT1p-GFP* | (CAI4) *RPS1/RPS1::pACT1-GFP* | (Barelle et al., 2004) |
| CAF2-dTomato | CAF2.1 (*Δura3::imm434/URA3*); pENO1-dTom-NATr | (Gratacap et al., 2013) |
| Eno1-iRFP | (CAI4) *NEUT5L/neut5l::FRT-ENO1p-iRFP-ADH1t-LoxP-*<br>*NAT1 RPS1/rps1::Clp10-URA3* | A.J.P Brown/D. Larcombe,<br>unpublished |
| 8880 (*rsr1Δ*) | (BWP17) *rsr1::ARG4/rsr1::HIS1 arg4::hisG/ARG4-*<br>*URA3::arg4::hisG* | (Hausauer et al., 2005) |
| JB6284/9955 (control<br>for *rsr1Δ*) | (BWP17) *his1::hisG/hisG::his1::HIS1 arg4::hisG/ARG4-*<br>*URA3::arg4::hisG* | (Bensen et al., 2002) |
| NGY355 (*pmr1Δ*) | (CAI4) *pmr1Δ::hisG/pmr1Δ::hisG, RPS1/rps1Δ::Clp10* | (Bates et al., 2005) |
| NGY356 *pmr1/PMR1* | (CAI4) *pmr1Δ::hisG/pmr1Δ::hisG, RPS1/rps1Δ::Clp10-*<br>*PMR1* | (Bates et al., 2005) |

### *A. dispar* microinjection

Needles were made using 1 mm OD borosilicate glass capillary (Harvard Apparatus) in a Narishige PC-10 micropipette puller at 60.4 °C. Needles were loaded with either *C. albicans* or PBS and placed into the manipulator before needle tips were removed using watchmaker’s forceps. An Eppendorf micromanipulator FemtoJet 4i was set to 400 injection pressure (pi) for 0.1 s with 20 hectopascals (hPa) Compensation pressure (Pc); the hold pressure was adjusted accordingly to counterbalance fluid uptake through the needle from capillary action during injection.

Molten 1% w/v agarose (Sigma-Aldrich) was poured into Petri dishes with hand-made moulds creating channels to hold the eggs. Eggs at 3 dpf were placed in these channels in ASW-filled dishes for 5-10 nl fungal inoculum injection into the ventral side of yolk sacs. Embryos were separated randomly into new petri dishes with ∼30 embryos containing fresh ASW and incubated as indicated. Water was changed daily, and dead or abnormal embryos terminated on inspection.

### Enumeration of fungal burden

To quantify fungal growth throughout infection, WT *A. dispar* embryos were injected into the yolk with approximately 50 cells per embryo of the desired *C. albicans* strain using injection mix of 5×10^6^ cells/ml (Table1) and incubated in ASW at 30 °C, 37 °C. At 0.5, 2, 6, 24, and 48 hpi, 3 embryos per group were randomly chosen and homogenised in penicillin-streptomycin solution 100 mg/ml (P4458, Sigma-Aldrich) using a hand-held motorised pestle mixer. Each embryo suspension was then plated on a separate YPD agar plate and incubated at 30°C for two days before counting the CFU. For the time points 24, 48 and 72 hpi, the homogenates were diluted with penicillin-streptomycin solution before plating.

### Automated imaging system and image analysis

To observe lethality, change in yolk diameter and fluorescence at regular intervals, an ACQUIFER Imaging Machine (Bruker) was used. *C. albicans*-injected, PBS-injected, and non-injected embryos were anaesthetised using a dilution of 1:9 of 0.4% tricaine methanesulfonate (MS-222) (Sigma) in 35 ppt ASW solution. One embryo per well was submerged in 170 μl of tricaine/ASW solution in a U-bottom 96 well plate and sealed with a plate sticker. A brightfield or fluorescent image was taken of an entire well using 2X or 4X objective, every 30 or 60 minutes for 72 hours. At each timepoint, 5 images were taken at the same z slice to observe heartbeat. A GFP channel (470nm) was used for ACT1p-GFP-infected *gch(-/-)* embryos. Embryos were incubated internally at 30, 37 and 40 °C.

The analysis was performed using the Fiji-software (Schindelin et al., 2012). For each well the Z-stack images were sequentially opened and stacked using *images to stack* option. The time point was noted when cardiac arrest occurred for each embryo. All the survival data were then noted in Excel sheet and transferred to GraphPad Prism version 10.0.3 (217) to generate a Kaplan-Meier survival graph (Goel et al., 2010).

### Antifungal treatment

*C. albicans* strain wild-type (SC5314) cells were collected, washed, and an injection mix of 5×10^6^ was prepared for the injection. Approximately 50 *Candida* cells were microinjected directly into the yolk of each *A. dispar* embryos. Control embryos received 5 to 10 nl Phosphate Buffered Saline (PBS). Fluconazole (FLC) was applied at different doses by directly exposing the embryos to artificial seawater containing the drug. A series of doses were used for the FLC agent. FLC was administrated at doses of 0, 2, 4, 8, 16 mg/L, respectively at 2 and 6 h post-infection at 37 °C. Survival rates of embryos were determined and plotted for different concentrations of drug. The fungal burden was assessed to evaluate the therapeutic efficiency of Fluconazole against different *C. albicans* strains.

### Dechorionation for fluorescent imaging

*A. dispar* embryos were rolled on fine sandpaper (p2000 grit size, waterproof) for 1 minute and then 3 were transferred to each Eppendorf tube containing 200 µl of 10 mg/ml pronase and 20 µl of 350 ppt ASW. Tubes were incubated at 30 °C or 37 °C for 1 hour with the temperature kept consistent thereafter in the incubator. Pronase was carefully removed, and eggs were briefly rinsed in ASW. Hatching enzyme (HE) was made by thoroughly homogenising ∼30 *A. dispar* embryos at 12 dpf with 200 µl ASW in Eppendorf tubes. Tubes were placed in a rotating disk for gentle mixing at 4 °C overnight before centrifuging at 13000 rpm at 4 °C for 10 mins. Supernatant (HE) was kept at -20 °C. Each tube had 400 µl HE added before incubation at 30 °C or 37 °C. Every hour, tubes were gently rotated. When chorions appeared disrupted, eggs were gently transferred using large-ended pipette tips to glass dishes with ASW for chorion removal using tweezers. Dechorionated eggs were transferred to a 24 well-plate, one per well, using a large-ended pipette tip, with 3 ml ASW for incubation at 30 or 37 °C overnight.

### Fluorescent microscopy

Images of *gch* (-/-) embryos infected with pACT1-GFP *C. albicans* strains were taken after dechorionation and embedding in agarose. An inverted fluorescent Olympus widefield microscope with 10x and 40x oil magnification objectives and an HBO lamp was used for embryos injected with RC-Yellow *C. albicans* strain, using both RFP and GFP filters.

A Delta Vision Elite High Resolution Microscope (Applied Precision) with a sCMOS_4.2 camera (Photometrics), running softWoRx 6.x was used for those injected with pACT1-GFP tagged *C. albicans*. Images were acquired using UPLFLN 40X/1.3NA oil immersion objective and GFP fluorescence was detected using a standard FITC filter. To image invasive hyphal growth, 43 optical sections were acquired at 0.7 µm increments and maximum intensity projections were generated using ImageJ.

Brightness and contrast of all images were adjusted, and appropriate colour added using ImageJ. Embryos were carefully dissected out of agarose before returning to a fresh well plate with ASW for incubation at 30 °C or 37 °C allowing embedding and imaging to be repeated.

### Statistical analyses

For differences in mean embryo survival time, the log rank test was employed using the Kaplan-Meier method. For differences in CFUs/embryo between temperatures, unpaired, two-tailed Student’s T tests were carried out for each time point post infection. p<0.05 was considered significant.

### Western blotting

Total protein samples from *A. dispar* embryos were prepared from five *A. dispar* embryos using 100 μl NuPAGE™ LDS Sample Buffer (4X) (Cat No. NP0008) containing 150 mM sodium chloride, 1.0% NP-40 (Triton X-100 can be substituted for NP-40), and 50 mM Tris pH 8.0, on ice for lysis and then homogenized by using a plastic pestle. Homogenised embryos were heated at 80-90 °C for 5 min and centrifuged at 14,000 rpm for 5 min. The supernatants were transferred to new 1.5 ml Eppendorf tubes and used immediately or stored at -20 °C until required. The amount of the supernatant which was loaded on a NuPAGE™ Mini Protein Gel (acrylamide 4%, ∼12% Bis-Tris pH7-7.5; Cat. No. NP0335BOX, Invitrogen) was set at 30 μl for embryo extracts and 7 μl of pre-stained protein standard ladder (SeeBlue® Plus2 Protein Standard, Thermo Fisher) in 1x NuPAGE MOPS running buffer at 120 V for 2 hours.

Transfer buffer 1x was used to transfer the proteins to the polyvinylidene difluoride (PVDF) membrane L16201177 (Bio-RAD) for 1 hour at 30 V, then its membranes were rinsed in ddH2O. Subsequently, the membrane was incubated in blocking solution (dried skimmed milk powder 5% w/v) in TBS for 2 hours and then probed with primary antibodies (Rabbit anti-mCherry 1:5000; Mouse anti-alpha Tubulin 1:1000) for overnight at 4 °C. Next day, the membrane was washed three times for 1 hour in 1x TBST (then incubated with the secondary antibodies (Goat-Anti Rabbit immunoglobulin HRP 1:1000; Goat-Anti Mouse immunoglobulins HRP 1:500) in blocking solution for 2 h at room temperature. To visualize signal development, the membrane was washed 3 times for 1 h with 1x TBST and incubated with a mixture of 1 ml luminal solution and 1 ml peroxide solution provided in the Immobilon Western HRP substrate kit (Thermo Scientific) for 2-5 minutes. The membrane was placed in an autoradiograph cassette with X-ray film for developing and the protein bands visualized.

## Acknowledgments

We extend our deepest gratitude to the ARC at the University of Exeter for their exceptional care and maintenance of the mutant and wild-type lines. We also extend our heartfelt appreciation to the Biological Sciences and MRC-CMM technical staff for their invaluable support and contributions to this research project.

## Supplementary Figures

**Fig. S1.**
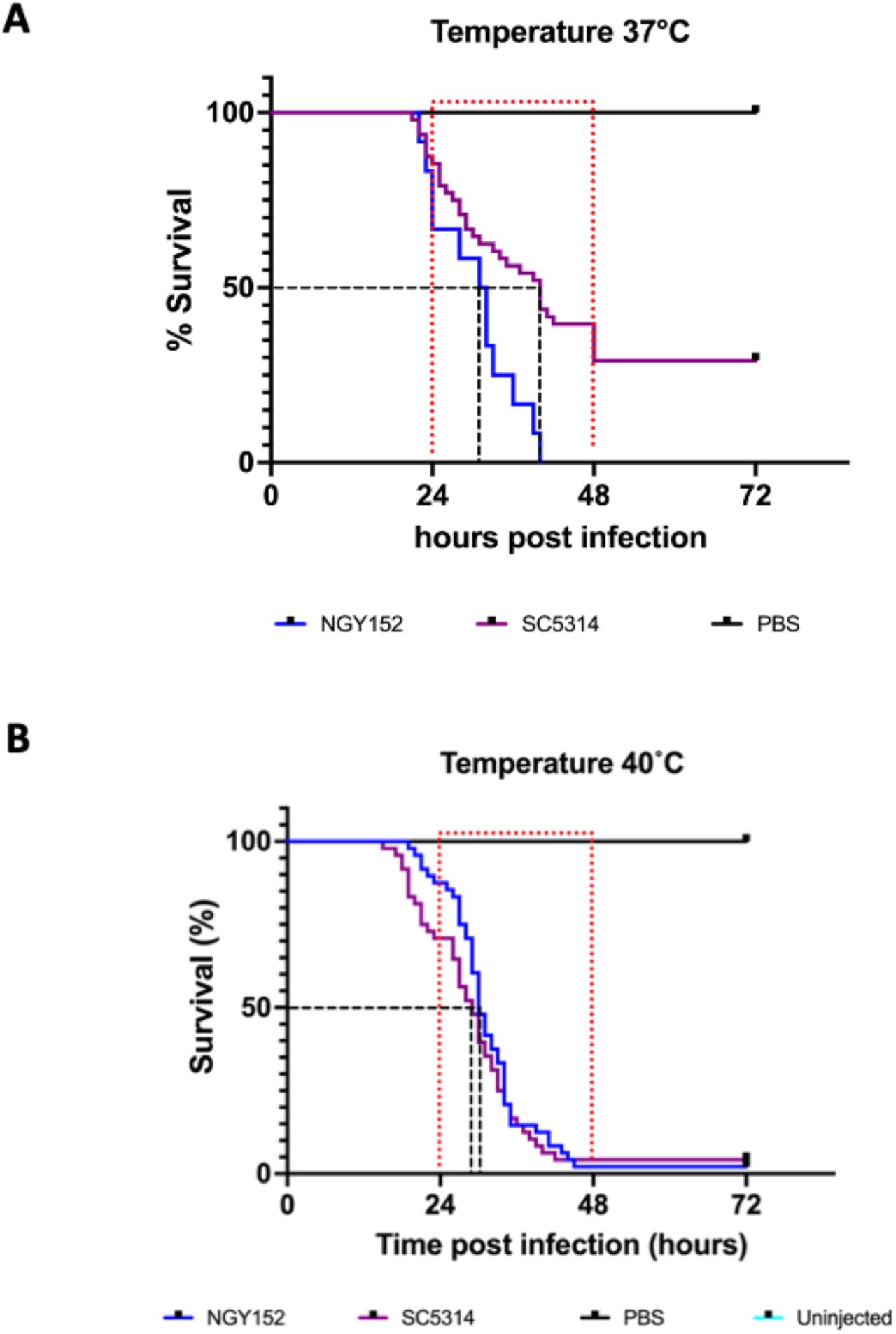
Comparative analysis of mortality time induced by NGY152 and SC5314 strains at A) body temperature and B) pyrexic temperature. NGY152 and SC5314, both exhibited peak embryo mortality in AKF embryos within the windows of 24 to 48 hours post *C. albicans* infection at human body temperature (37 °C). Statistically significant difference (log-rank (Mantel-Cox) test p-value < 0.0001) was observed between the control and pyrexic conditions (40 °C). The experiment conducted at pyrexic temperatures had n = 4 replicates, with 12 embryos per condition.

**Fig. S2.**
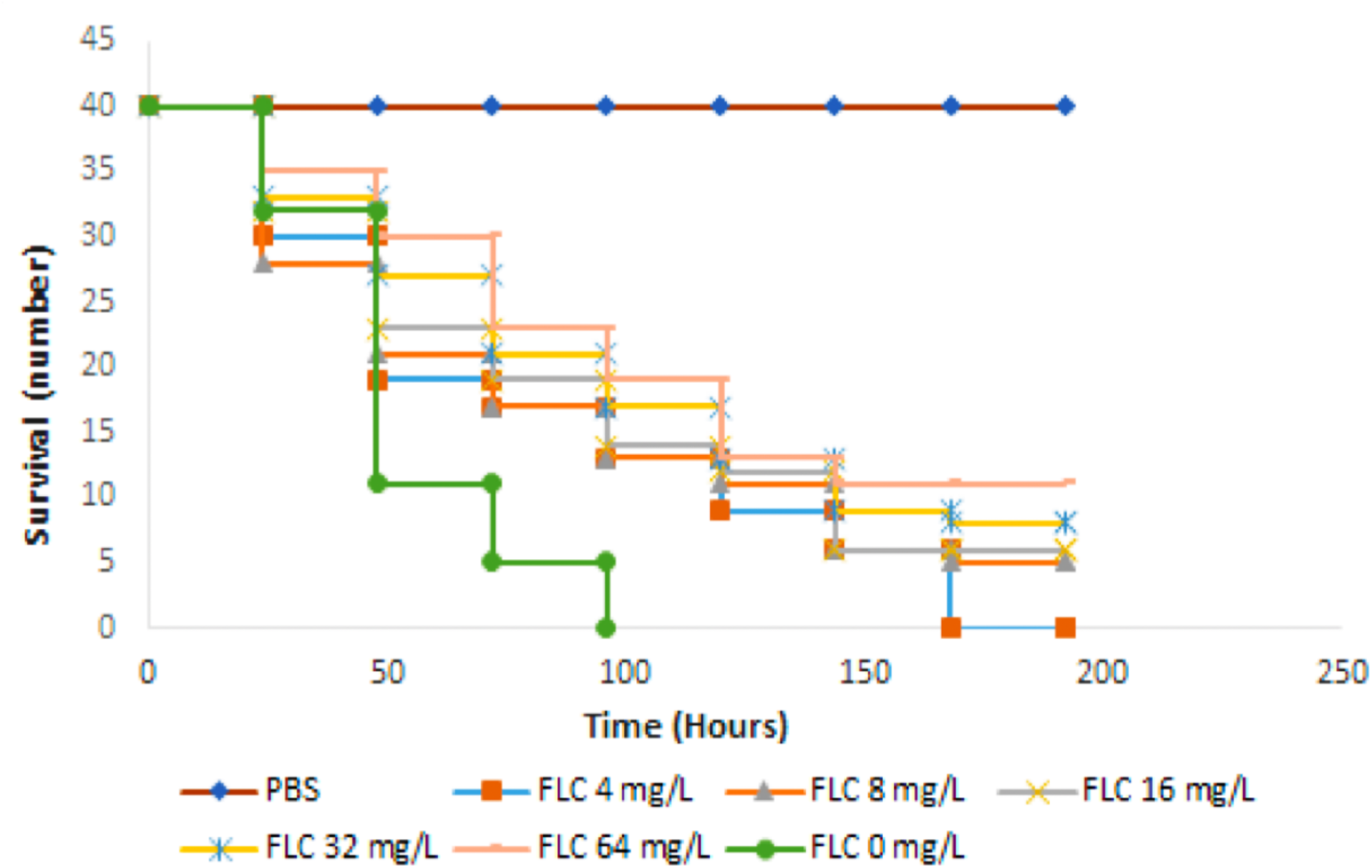
Delayed antifungal treatment shows reduced effectiveness against C. albicans-colonised A. dispar embryos. (A) *A. dispar* embryos at 3 days post fertilization were infected with 5 x 10^6^ cells/ml *C. albicans (strain name).* At 6 h post infection, embryos were treated with 4, 8, 16, 32, or 64 mg/L fluconazole (FLC). Although treatment delayed mortality by ∼ 3d, this was not dose-dependent (at the concentrations used). Control groups, including PBS-injected embryos, showed 100% survival while infected but untreated embryos showed rapid mortality.

**Fig. S3.**
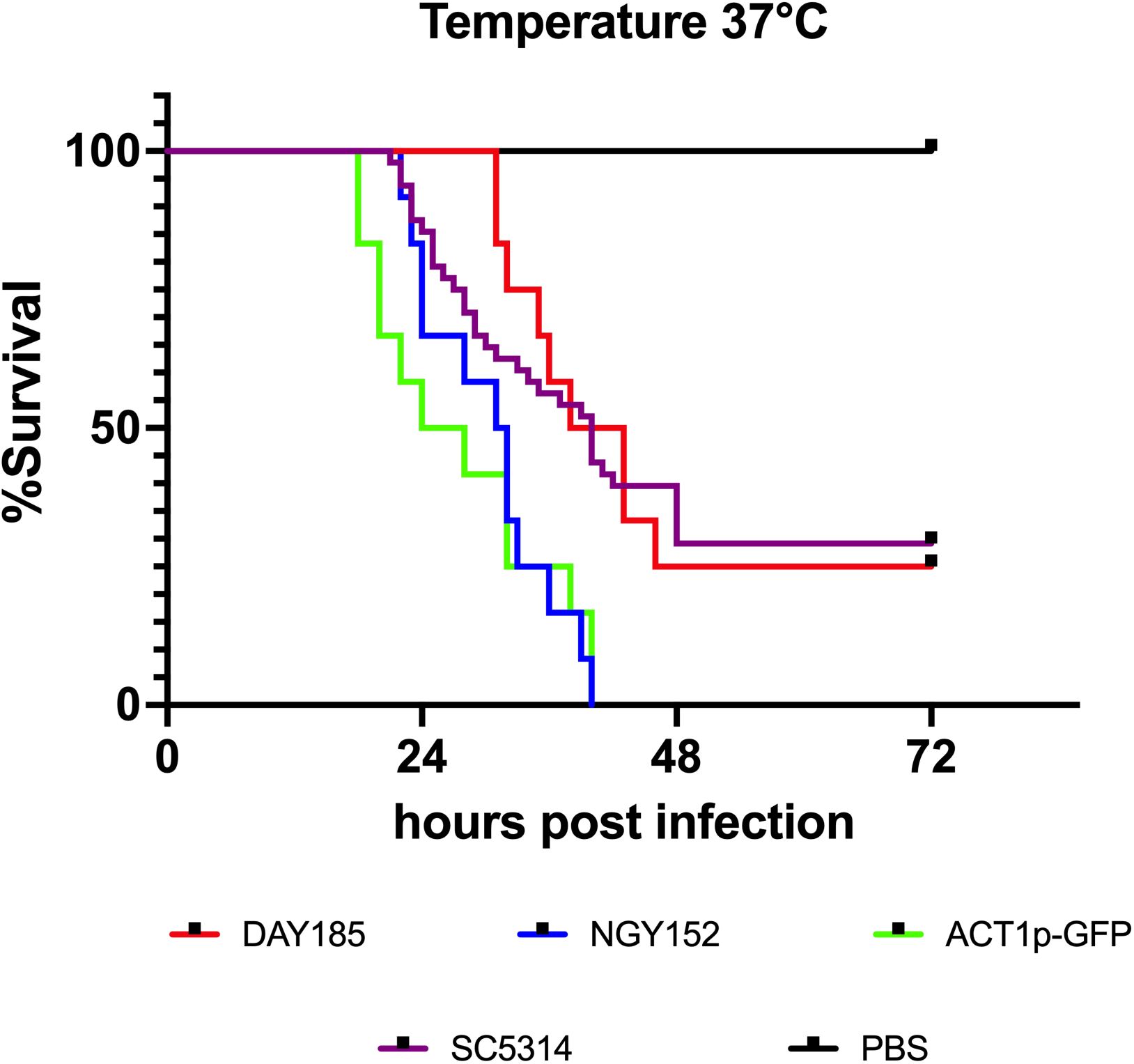
Comparative analysis of mortality time induced by different *C. albicans* strain at body temperature. Day185 and NGY 152, ACT1p-GFP and SC5314, all exhibited peak embryo mortality in AKF embryos within 48 hours post-*C. albicans* infection, with over 75% of embryos showing signs of lethality during this period (Log-rank (Mantel-Cox) p-value <0.0001).

**Fig. S4.**
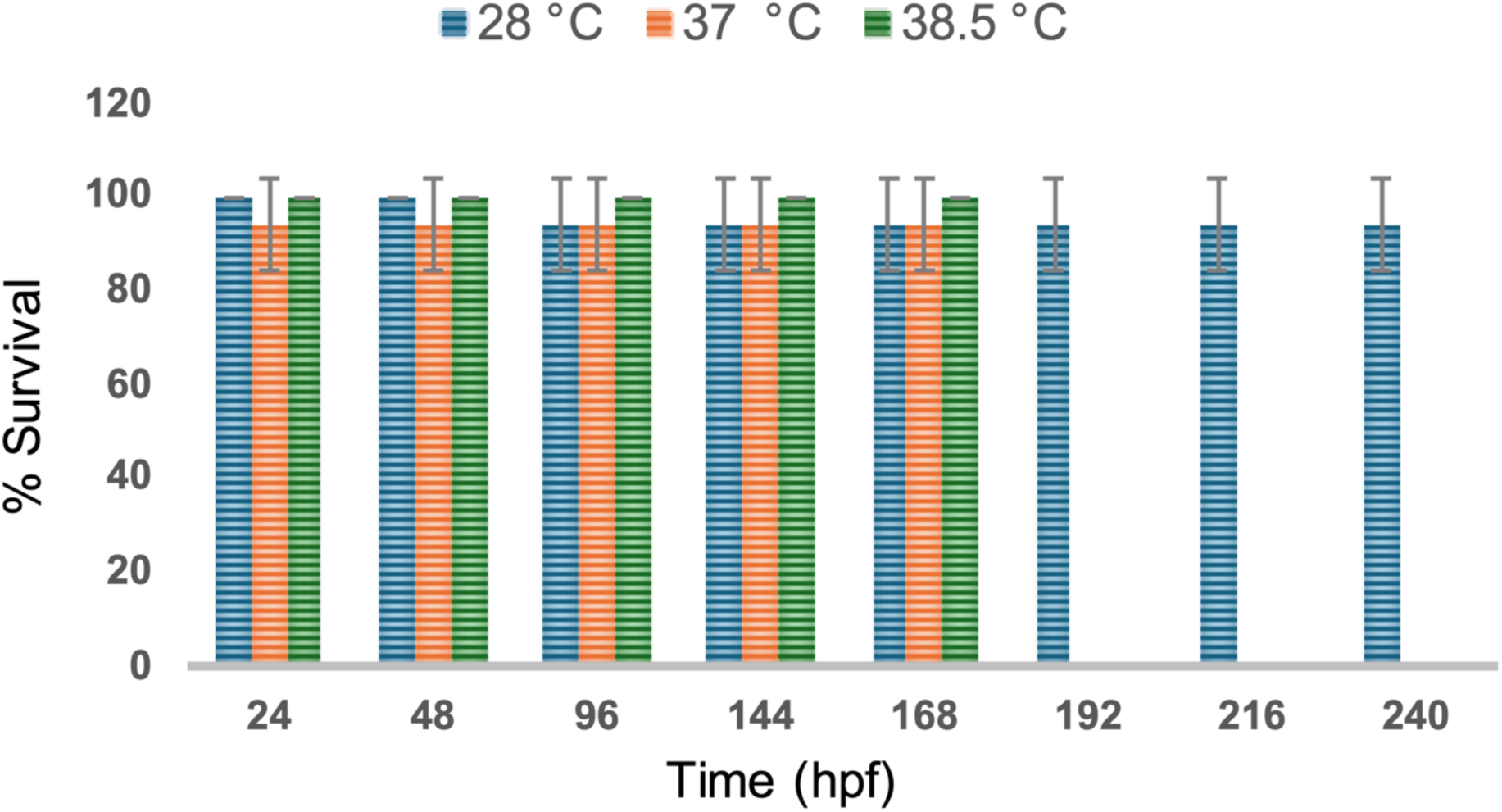
AKF embryos exhibit accelerated development at 37°C and 38.5 °C without compromising viability. AKF embryos were immediately transferred to a 37°C incubator upon collection from breeding chambers and monitored daily for recording any mortality. The experiment was conducted with three biological replicates (N=3), each consisting of 7 embryos per petri dish. Embryos incubated at 37 and 38.5 °C hatched at 168 hpf.

